# Neurons in human pre-supplementary motor area encode key computations for value-based choice

**DOI:** 10.1101/2021.10.27.466000

**Authors:** Tomas G. Aquino, Jeffrey Cockburn, Adam N. Mamelak, Ueli Rutishauser, John P. O’Doherty

## Abstract

Adaptive behavior in real-world environments demands that choices integrate over several variables, including the novelty of the options under consideration, their expected value, and uncertainty in value estimation. We recorded neurons from the human pre-supplementary motor area (preSMA), ventromedial prefrontal cortex (vmPFC) and dorsal anterior cingulate to probe how integration over decision variables occurs during decision-making. In contrast to the other areas, preSMA neurons not only represented separate pre-decision variables for each choice option, but also encoded an integrated utility signal and, subsequently, the decision itself. Conversely, post-decision related encoding of variables for the chosen option was more widely distributed and especially prominent in vmPFC. Our findings position the human preSMA as central to the implementation of value-based decisions.

## Introduction

Humans and other animals can make decisions in a manner that maximizes the chance of obtaining rewards. Computational theories of decision-making suggest that doing so relies on a number of variables.^1^ Most studied among these is the expected value (EV) associated with an option. By comparing options with varying EVs, it is possible to guide behavior toward higher expected future reward. However, in the real-world, the relationship between actions and their subsequent outcomes is often uncertain, and as such, one needs to consider not only the expected reward, but also its estimation uncertainty.^2, 3^ Another relevant feature is the novelty of an option – novel options can potentially provide new opportunities to gain reward.^4^ These features can be utilized to resolve an often encountered dilemma in decision-making: whether to explore uncertain options that could yield richer reward, or to exploit an option with known rewards.^5, 6^

How does the human brain represent the decision variables associated with the available options and how are they integrated to make a decision? One possibility is that neurons encode a utility signal that integrates over relevant decision variables for a given option and that this integrated utility is then used as an input to the decision process. Alternatively, these variables could be encoded in non-overlapping neuronal populations and be integrated at the population level to inform action selection.

Studies in rodents and non-human primates have reported neurons throughout the prefrontal cortex that correlate with EV,^7–12^ uncertainty^13–16^ and novelty.^17–19^ Most human studies have been restricted to non-invasive methods such as functional magnetic resonance imaging (fMRI), revealing roles in value-based decision making for the vmPFC,^20–23^ dorsal anterior cingulate cortex (dACC),^24^ and preSMA.^21, 25^ Overall, these areas encode decision variables such as EV,^9, 20, 21, 23, 25^ uncertainty,^23, 26, 27^ and outcomes,^9, 28^ while novelty related effects have also been found in the dopaminergic midbrain and striatum.^4, 29–31^ Some studies reported value computations in prefrontal cortex utilizing intracranial EEG (iEEG) from depth and grid electrodes.^32, 33^ While this approach affords greater temporal resolution than fMRI, iEEG reflects pooled activity across large numbers of neurons with a similar lack of spatial selectivity as fMRI.

We overcome these limitations by recording from single neurons in preSMA, dACC, and vmPFC while human patients with drug resistant epilepsy undergoing invasive electrophysiological monitoring performed a decision-making task specifically designed to dissociate EV, uncertainty and novelty. We sought to determine how neurons in these three brain areas differ during decision-related computations, to address whether these variables are integrated into a utility signal at the level of single neurons, and to probe how these signals might be utilized for informing choice. Additionally, we aimed to distinguish neurons that encode stimulus features and choice from those that evaluate the consequences of the decision. Finally, we could identify neurons encoding outcomes and prediction errors, to ascertain how these regions contribute to updating decision information following feedback at the neuronal level. Thus, this study afforded us an unparalleled opportunity to investigate the role of human prefrontal neurons across multiple stages of value-based decision-making: from the representation of individual decision variables, through to integration of these variables, up to choice and ultimately feedback.

## Results

### Task and behavior

We recorded 191 vmPFC, 137 preSMA and 108 dACC single neurons (436 total) in 22 sessions from 20 patients chronically implanted with hybrid macro/micro electrodes for epilepsy monitoring (Fig. 1 A). Patients performed a two-armed bandit task^34^ designed to separate the influence of EV, uncertainty and novelty on decision making, divided into 20 blocks consisting of 15 binary choices. On each trial, participants used a button box to choose between two uniquely identifiable bandits presented on the left or the right of the screen (Fig. 1 B). Following a time delay, a feedback screen then announced the binary outcome (win/no win). The experimental design included two critical features. Firstly, participants were informed that the probability of each bandit delivering a reward was fixed for the duration of each block, but randomized across blocks. Secondly, both novel and familiar stimuli were systematically incorporated into the set from which options could be drawn during a block, resulting in pairs of bandits that varied in terms of EV, uncertainty and novelty.

**Figure 1:**
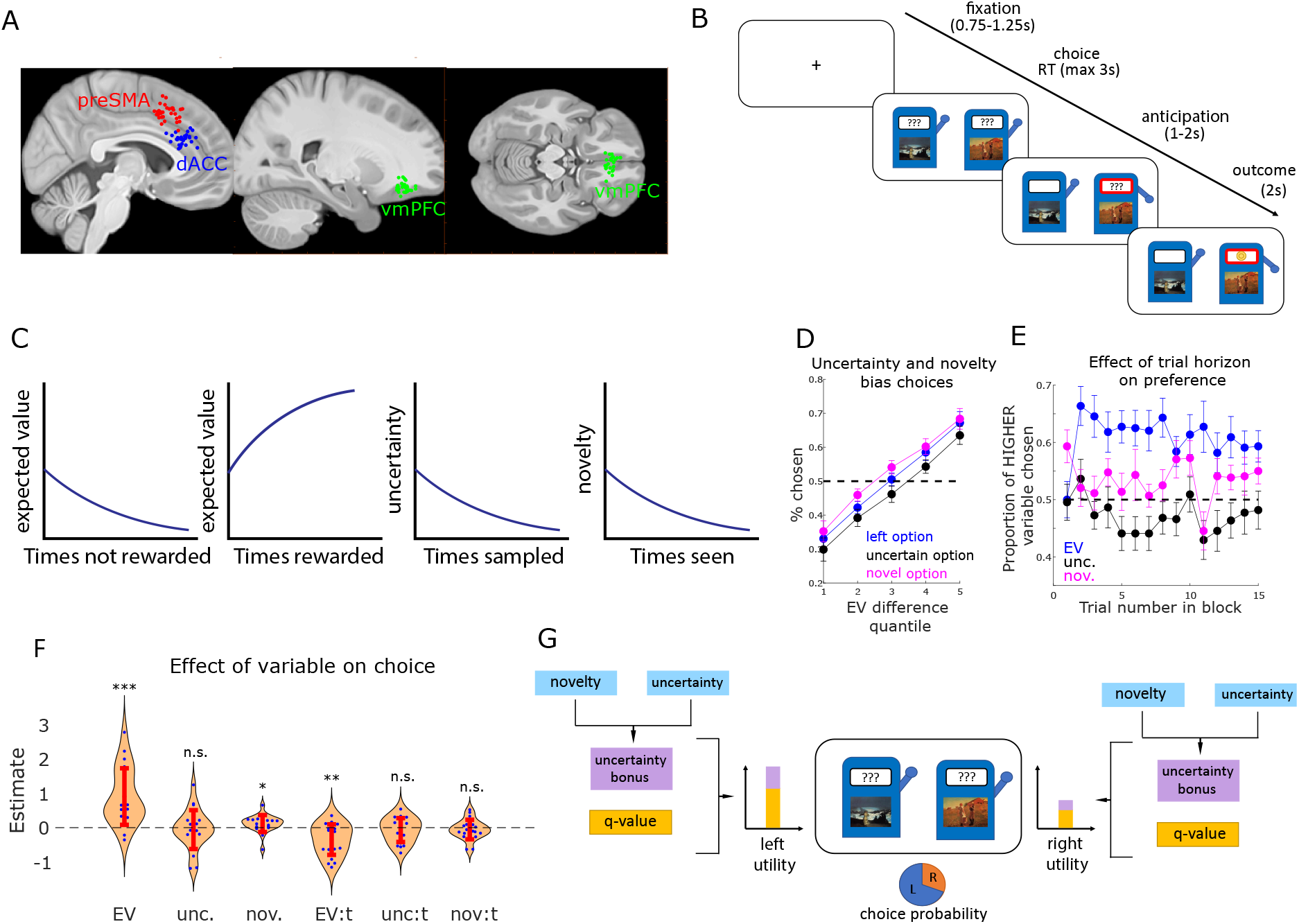
Exploration task, electrode positioning and behavior. (a) Electrode positioning. Each dot indicates one electrode in one patient in preSMA (red), dACC (blue), or vmPFC (green). (b) Trials were structured in fixation, decision, anticipation and feedback stages. (c) Schematic indicating how q-values, uncertainty and novelty of stimuli vary as a function of the past history of rewards, sampling and exposures. (d) Expected value correlates with choice, biased by novelty and uncertainty. Patients chose the left option (blue), the more uncertain option (black), or the more novel option (magenta) as a function of chosen minus unchosen expected value. (e) Proportion of trials in which patients chose the option with higher expected value (blue), uncertainty (black), or novelty (magenta), as a function of trial number. Dots and bars indicated means and standard errors, respectively. (f) Logistic regression coefficients for expected value, uncertainty, novelty, and interactions with trial number. Dots and bars indicate fits for each patient and standard error, respectively (* = p<0.05; ** = p<0.01; *** = p<0.001, t-test). Positive values indicate seeking behavior. (g) fmUCB model. Novelty and uncertainty generate an uncertainty bonus, which composes utility along with q-values.

We first assess how EV, uncertainty and novelty related to behavior. Within each block of trials, EV and uncertainty were quantified as the average proportion of wins and total number of samples within a given block of trials respectively, while novelty was defined as the total number of times a stimulus had been shown across the entire experiment (see Fig. 1 C). Uncertainty and novelty biased value-based decisions in distinct directions: while on average patients preferred options with higher EVs over options with lower EVs (*p* < 10^−51^, *T* = 18.2, linear regression), they sought them more often if they were also the more novel option, than if they were also the more uncertain option (*p* < 0.01, *T* = 2.73, two-sided t-test) (Fig. 1 D). This was not the result of changing preferences over time because trial number did not correlate with how often patients sought the option with higher uncertainty (*p* = 0.31, *T* = −1.00, linear regression) or higher novelty (*p* = 0.76, *T* = −0.29, linear regression) (Fig. 1 E). We then used a logistic regression to correlate decision variables and choices (see Materials and Methods section for details on logistic regression analysis), with EVs, uncertainty, and novelty as predictors. Model coefficients (Fig. 1 F) indicated that patients were EV-seeking (*p* < 10^−4^, *T* = 5.15, t-test) and novelty seeking (*p* < 0.05, *T* = 2.26, t-test), with a negative effect of the interaction between EV and trial number (*p* < 0.01, *T* = −3.67, t-test), suggesting a deviation from optimal outcome integration. Importantly, we also confirmed that patient behavior reflected value reset at the start of each block (see supplement), indicating that patients understood the task structure and learned how to choose the more advantageous options from their past experiences. Additionally, novelty and uncertainty correlated with behavior in a separable manner: while patients tended to be novelty-seeking overall, some patients avoided uncertainty whereas others were uncertainty seeking.

We compared two candidate computational models for explaining patient behavior (see Supplementary Material for details on model comparison). The first model (familiarity modulated upper confidence bound (fmUCB), Fig. 1 G) was developed to describe behavior and fMRI data in a neurotypical population performing this same task.^34^ In this model, novelty promotes optimistic value initiation and modulates uncertainty to form an uncertainty bonus, which is added to q-values to construct stimulus utilities. The second model (linear novelty) relied on a linear combination between q-values, uncertainty and novelty to construct utilities. Using hierarchical Bayesian inference on patients’ behavioral data^35^ we determined the fmUCB model to explain behavior best, and a posterior predictive check showed that there are no systematic discrepancies between behavioral and simulated data. These behavioral modeling results show that a familiarity gating mechanism is appropriate to model the behavior of our participants. For subsequent neural data analyses we therefore used the variables derived from the fmUCB model as regressors.

### Representation of separate stimulus features for individual options

We next probed the neural representation of stimulus features by examining whether the q-value, uncertainty, or novelty of each option presented on the screen was represented by neurons in our regions of interest using a Poisson GLM, with these features as regressors (for a complete list of encoding models, see Table S2). As these variables pertain to each stimulus being considered on a given trial and are not contingent on the choice of option that is subsequently made, they are candidate variables for acting as an input to the decision process.

We then grouped neurons according to their sensitivity to features associated with the left or right option, which we refer to as positional q-value, uncertainty bonus, and novelty neurons respectively (see Fig. 2 D for an example). To determine whether activity in a brain area significantly correlated with these positional stimulus features, we tested whether the selected number of neurons were larger than expected by chance (Figs. 2 A-C). All subsequent neuron count results were Bonferroni corrected for the number of time windows and brain areas in which we tested for a significant neuron count.

**Figure 2:**
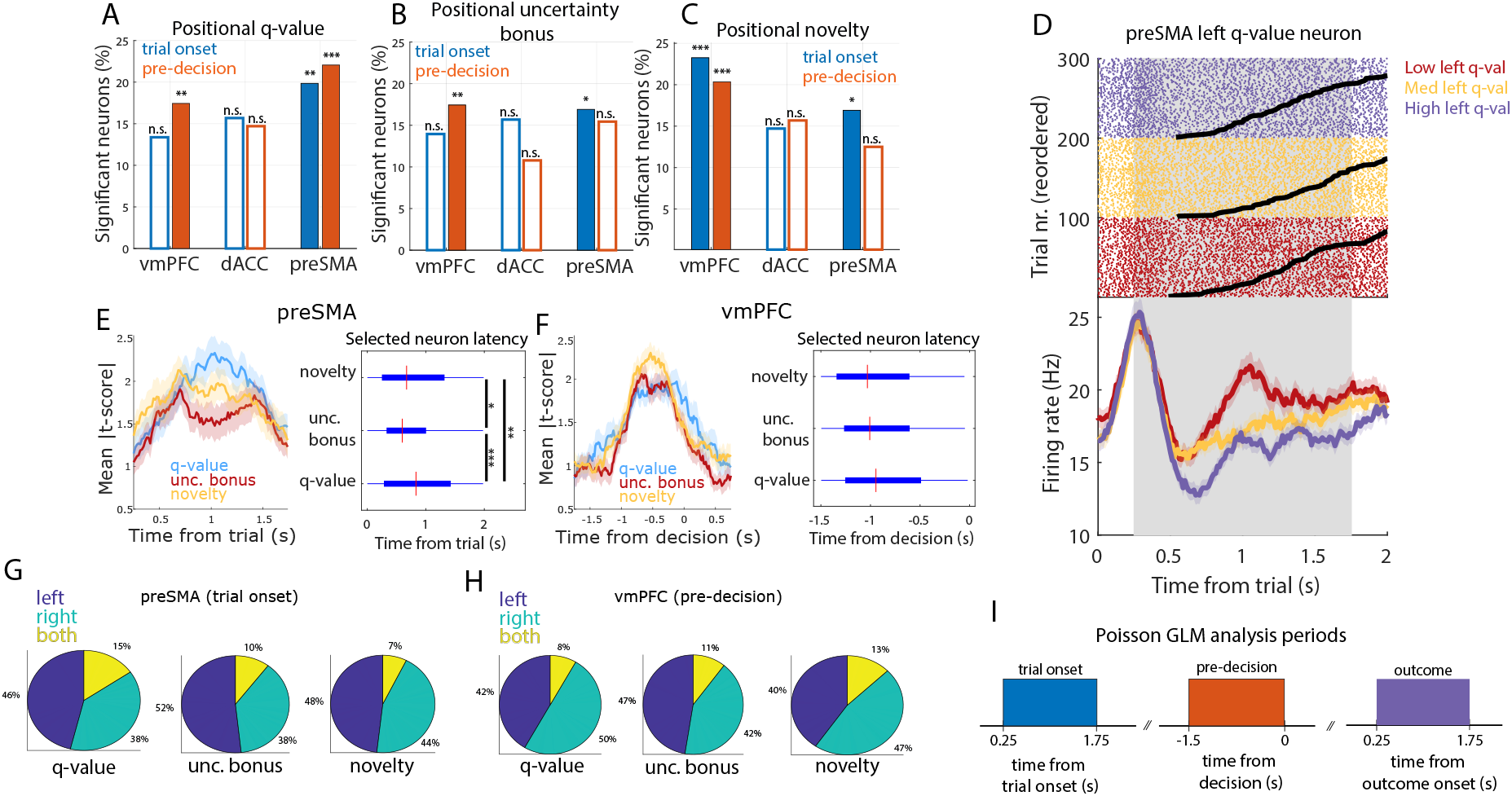
Encoding positional utility components in preSMA and vmPFC. (a) Percentage of neurons sensitive to positional q-value, in the trial onset (blue) and pre-decision (orange) periods (* = p<0.05; ** = p<0.01; *** = p<0.001, binomial test). Hollow bars indicate non-significant counts. (b) Same, for positional uncertainty. (c) Same, for positional novelty. (d) Left q-value preSMA neuron. Top: Trial aligned spike raster plots. Black lines indicate the response time. For plotting, we sorted trials by q-value tertile (purple: high; yellow: medium; red: low). Bottom: Trial onset aligned PSTH (bin size = 0.2s, step size = 0.0625s). Shaded areas indicate standard error. (e) Selected preSMA neuron timing in the trial onset period. Left: Mean absolute t-score from the Poisson GLM analysis, in q-value (blue), uncertainty bonus (red), and novelty (yellow) neurons. Shaded areas indicate standard error. Right: Box plots of latency time across trials for all q-value, uncertainty bonus, or novelty neurons (* = p<0.05; ** = p<0.01; *** = p<0.001, two-sided rank-sum test). (f) Same, for vmPFC neurons in the pre-decision period. (g) Proportion of preSMA sensitive neurons encoding positional q-values (left), uncertainty bonuses (center), or novelty (right), for one or both options. (h) Same, for vmPFC neurons in the pre-decision period. (i) Time windows for all analyses (trial onset, pre-decision, and outcome).

This analysis revealed prominent encoding of positional q-value, uncertainty bonus, and novelty during the trial onset period (19.9%, *p* < 0.01/6; 16.9%, *p* < 0.05/6; and 16.9%, *p* < 0.05/6, respectively, binomial test) and robust encoding of positional q-value during the pre-decision period in the preSMA (22.1%, *p* < 0.001/6, binomial test). In contrast, neurons in the vmPFC encoded positional q-value and uncertainty during the later pre-decision period, (17.4%, *p* < 0.01/6; 17.4%, *p* < 0.01/6, respectively), whereas novelty was encoded during both time periods (trial onset: 23.2%, *p* << 10^−5^/6; pre-decision: 20.3%, *p* < 0.001/6, binomial test). None of the selected cell counts were significant in dACC (Figs. 2 A-C). This indicates that preSMA and vmPFC neurons encode the variables which can serve as input to the decision process, aligned to trial onset in preSMA, and to decision in vmPFC.

Given the prominent role of preSMA and vmPFC in encoding all components of value in the trial onset period and the pre-decision period respectively, we investigated the temporal activity patterns for the selected neurons in these two areas. We repeated the Poisson GLM analysis described above in sliding time windows for visualization (Figs. 2 E-F) and performed a Poisson latency analysis in neurons which were exclusively sensitive to one of the tested variables^36^ to compare onset latencies (see Materials and Methods). In preSMA (Fig. 2 E), positional uncertainty neurons (median time: 0.59s) became active first, followed by positional novelty neurons (median time: 0.67s, *p* < 0.05, two-sided rank sum test) and positional q-value neurons (median time: 0.83s, *p* < 10^−6^, two-sided rank sum test). Positional novelty neurons were found to be active significantly earlier than positional q-value neurons (*p* < 0.01, rank sum test). In vmPFC, median activation times relative to the time of decision were not significantly different for any of the selected sub-populations. Therefore, neurons in preSMA, but not in vmPFC, which are sensitive to distinct value components, have significantly different latencies. Notably, preSMA q-value neurons have longer latencies than novelty and uncertainty bonus neurons.

Neurons coding for “positional” components of value were predominantly sensitive exclusively to a single option: for every variable, less than 15% of selected neurons in preSMA (trial onset) and in vmPFC (pre-decision) were found to be sensitive to features of both the left and right options available (Figs. 2 G-H). This indicates that these neural sub-populations encode q-values, uncertainty and novelty preferentially for individual stimuli instead of a total or averaged signal across options, which would be a possible interpretation if there had been prominent simultaneous encoding of both left and right options.

For completeness, we repeated the above analysis using values for q-value, uncertainty bonus and novelty as derived from the alternative linear novelty behavioral model. This revealed qualitatively similar results for positional q-value and novelty encoding. However, no significant neuron counts for positional uncertainty bonus encoding was found in any brain area. This highlights how the fmUCB model is specifically capable of describing uncertainty bonus representations accounting for neuronal activity.

Taken together, these findings suggest that stimulus features for the two available options are first encoded in the preSMA (trial onset period), followed by the vmPFC in the pre-decision period. This encoding was in the form of most neurons signaling stimulus features for one but not both options, which would be expected from a signal that serves as an input to the decision process.

### Single neuron encoding of integrated utility for individual stimuli

To determine whether neurons represented an integrated utility for each decision option (incorporating EV, uncertainty and novelty), we used the utility signal derived from our fmUCB model. We performed a Poisson GLM encoding analysis (Fig. 3 A,B) with left utility, right utility and decision as regressors. We found that a significant number of preSMA neurons encoded left utility after the trial onset (16.2%, *p* < 10^−5^/6, binomial test), and in the pre-decision period (13.2%, *p* < 0.001/6, binomial test). Similarly, a significant number of preSMA neurons encoded right utility after trial onset (11.8%, *p* < 0.01/6, binomial test), and in the pre-decision period (11.8%, *p* < 0.01/6, binomial test). This result suggests that single neurons in the preSMA encode an integrated utility signal for individual choice options. Alternatively, it is possible that neurons correlating with utility in our regression analysis are mostly reflecting the effects of q-value per se.

**Figure 3:**
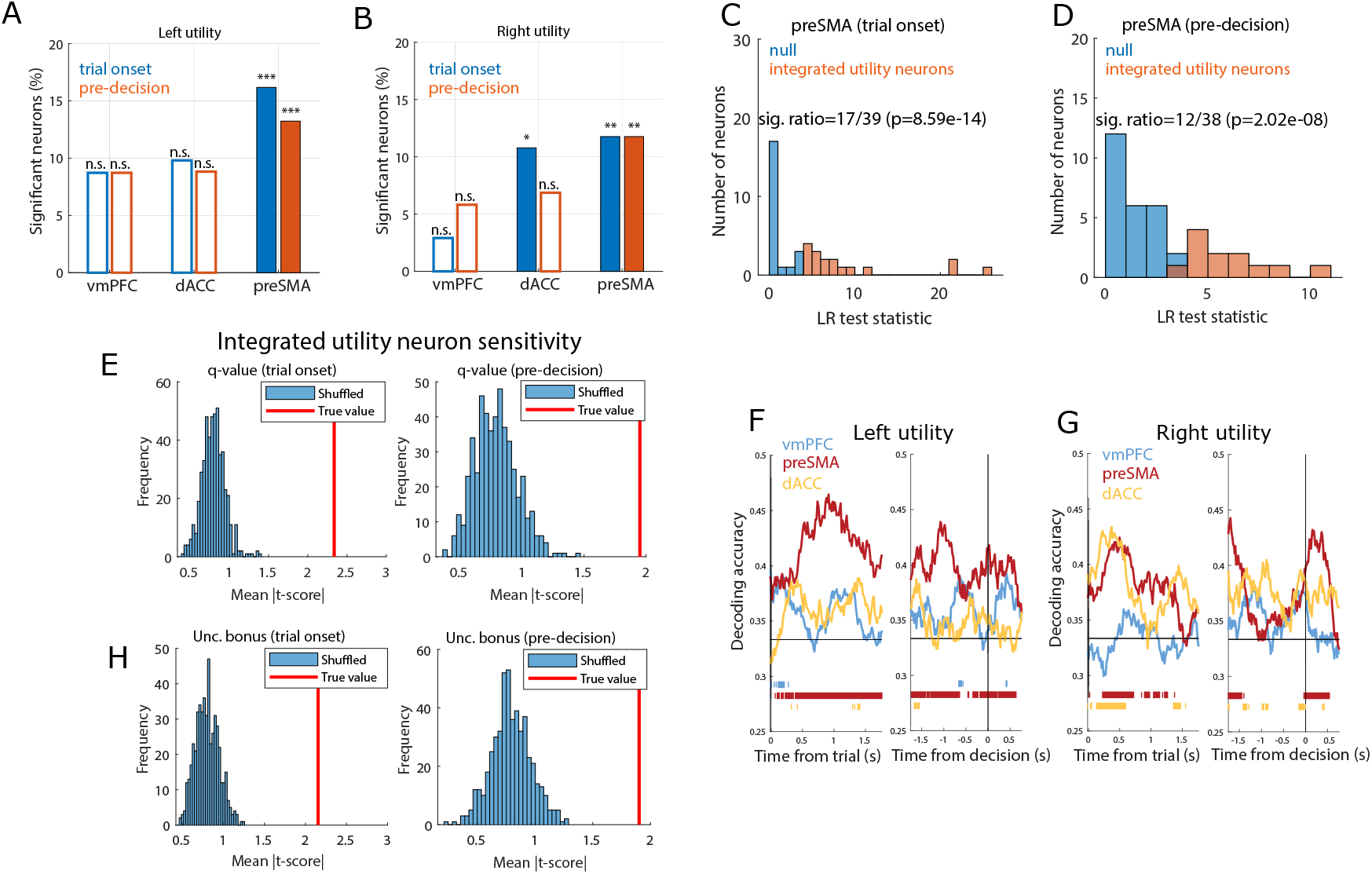
Neurons in preSMA encode integrated utility. (a) Percentage of left stimulus utility neurons in vmPFC, dACC, and preSMA, in the trial onset (blue) and pre-decision (orange) periods (* = p<0.05; ** = p<0.01; *** = p<0.001, binomial test). Hollow bars indicate non-significant counts. (b) Same, for right stimulus utility. (c) Likelihood ratio test statistics across candidate preSMA integrated positional utility neurons, in the trial onset period. Neurons whose activity was better explained by a model containing q-values and uncertainty bonuses were classified as integrated utility neurons (orange). For the remaining ones (blue) the null model restricted to q-values was not rejected. (d) Same, for pre-decision period. (e) Integrated utility preSMA neuron sensitivity to q-values. Red lines indicate the mean absolute t-score across integrated utility neurons. Histograms include mean absolute t-scores for 500 iterations of bootstrapped null models with shuffled firing rates. Left: trial onset period; Right: pre-decision period. (f) dPCA population decoding performance for left utility for vmPFC (blue), preSMA (red), and dACC (yellow). Bars indicate periods of time where decoding accuracy was significantly above chance. Left: trial onset period; Right: pre-decision period. (g) Same, for right utility. (h) Same as (e), for uncertainty bonuses.

To test this hypothesis, we defined the sub-populations of preSMA neurons identified either as q-value or utility neurons as candidate neurons for an integrated utility signal. To determine whether they encoded an integrated utility signal versus q-value alone, we performed a likelihood ratio test (*p* < 0.05) comparing the performance of a model containing q-value, uncertainty, and decision regressors versus a null restricted model containing only q-value and decision (see Materials and Methods), while predicting each candidate neuron’s spike count. The null restricted model was rejected for 44% (17/39) of preSMA candidate neurons at trial onset and for 32% (12/38) of preSMA candidate neurons in the pre-decision window (Figs. 3 C-D). Therefore, a significant portion of candidate neurons in preSMA qualified as integrated utility neurons (trial onset: *p* < 10^−13^; pre-decision: *p* < 10^−7^). These integrated utility neurons collectively encoded the main components of utility (q-values and uncertainty bonuses) at a higher level than expected by chance (*p* < 0.002 in all instances, permutation test), further confirming their role in computing an integrated signal (Figs. 3 E,H).

### Population decoding of integrated utility as an input to the decision process

Building upon these results demonstrating that single neurons in preSMA and vmPFC encode stimulus features that could support the decision making process, we next tested when and where it was possible to decode an integrated stimulus utility value from neural populations. To do so, we consider the firing patterns of all neurons from each brain region across all trials, employing demixed principal component analysis^37^ (dPCA) to reduce the data dimensionality (see Materials and Methods).

We performed two separate analyses for left and right utilities, including the decision itself (i.e. left vs right choice) as a marginalization in both analyses (Figs. 3 F-G). Left and right option utility was decodable in preSMA, both following trial onset and preceding the button press (Figs. 3 F-G, significant time periods are indicated in the figure). Compatible with the cell selection results, neither left nor right side utility was robustly decodable from vmPFC.

Thus, these results suggest that preSMA encodes an integrated utility signal that encompasses both q-values and uncertainty. At the population level, the utility for each decision option was decodable in preSMA even after demixing utility from the decision, indicating that the utility for each of the two possible decision options is represented at the population level. Together, these findings suggest that preSMA neurons represent the signals needed as an input to the decision making process.

### Decision is represented later than stimulus utility

At the level of single neurons, the decision was encoded in the preSMA only in the pre-decision period (Fig. 4 A), in which 14.0% (*p* < 10^−4^/6, binomial test) of neurons were decision selective (Fig. 4 F shows an example). In preSMA, neurons that encoded left or right utility were largely distinct from those encoding the decision (Fig. 4 C), with no significant overlap for either (*p* = 0.22 and *p* = 0.28, Jaccard test, see Materials and Methods). Neither at the single-neuron (Fig. 4 A) nor the population level was the decision represented in vmPFC or dACC (Fig. 4 D), indicating a privileged role for the preSMA in representing choices in our task. We therefore restrict the following analysis to the preSMA.

**Figure 4:**
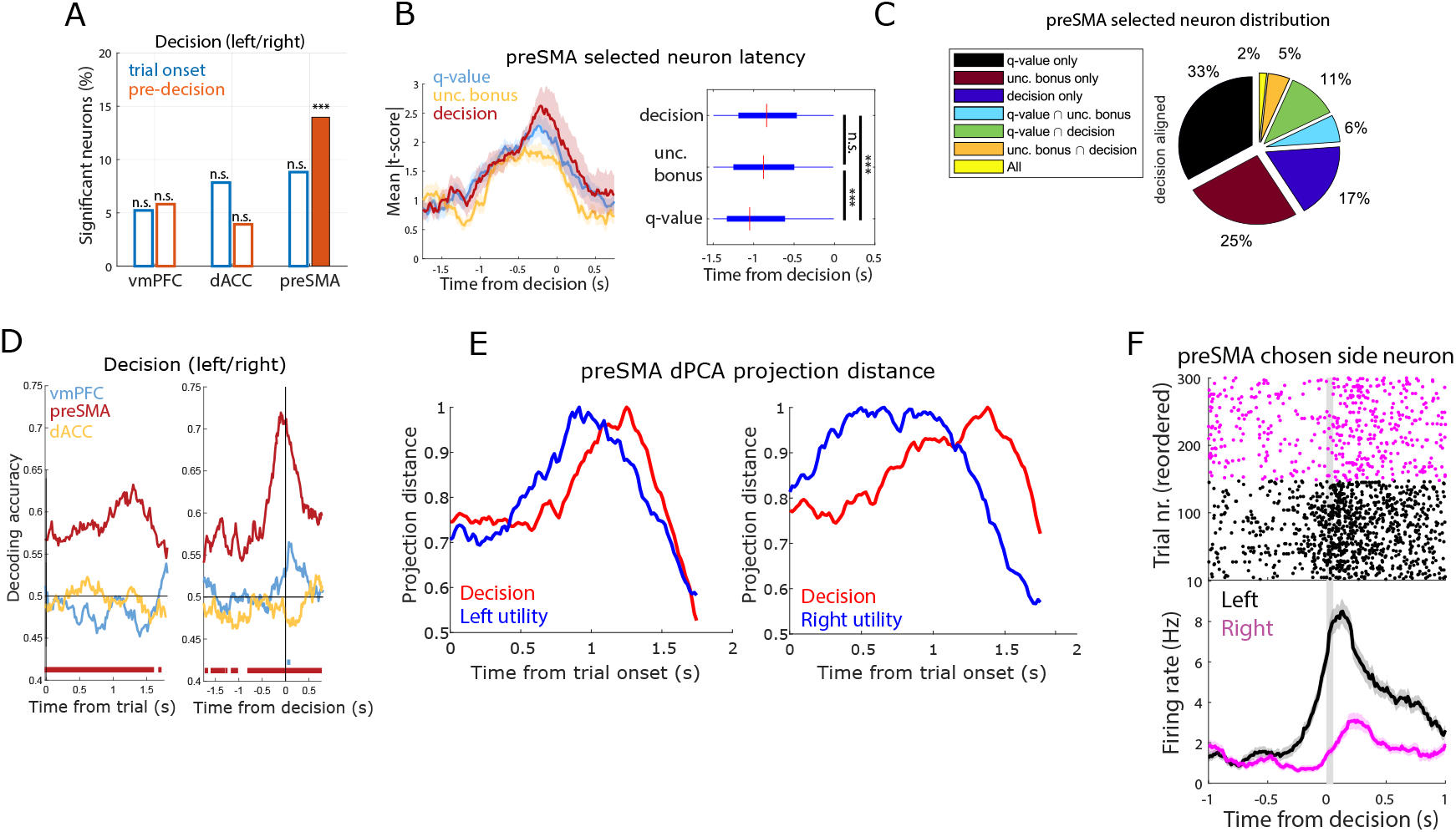
PreSMA encodes decisions. (a) Percentage of decision neurons (left vs. right choice) in vmPFC, dACC, and preSMA, in the trial onset (blue) and pre-decision (orange) periods (*** = p<0.001, binomial test). Hollow bars indicate non-significant counts. (b) Sensitive preSMA neuron timing in the pre-decision period. Left: Mean absolute t-score for q-value (blue), uncertainty bonus (yellow), and decision (red). Shaded areas indicate standard error. Right: Latency time box plots for all q-value, uncertainty bonus, or decision neurons (*** = p<0.001, two-sided rank-sum test). (c) Proportion of preSMA neurons (pre-decision period) for q-value, uncertainty bonus, decision and combinations thereof. (d) dPCA decision decoding for vmPFC (blue), preSMA (red), and dACC (yellow). Bars indicate significant times, comparing to a bootstrapped null distribution. Left: trial onset period; Right: pre-decision period. (e) Normalized Euclidean distance between dPCA projections onto principal utility components (blue), between low and high utility trials, and decision components (red), between left and right decision trials, with left (left) or right (right) utility marginalizations. (f) Example preSMA decision neuron. Top: Raster plot. For plotting, we sorted trials in left (black) and right (magenta) decisions. Bottom: PSTH (bin size = 0.2 s, step size = 0.0625 s). Gray bar indicates button press. Shaded areas indicate standard error.

Relative to the time of response, a single-unit analysis showed that q-value neurons responded first at −1.04*s*, earlier than uncertainty bonus neurons at −0.87*s* (*p* < 10^−3^, two-sided rank-sum test), and decision neurons at −0.83*s* (*p* < 10^−3^, two-sided rank-sum test). At the population level, we projected neural data onto the dPCA demixed principal components components separately for low/high utility trials, and for left/right decisions. We then examined the Euclidean distances between these trajectories as a function of time. This revealed that the distance in state space was maximal for positional utility earlier than for decisions (Fig. 4 E). Relative to trial onset, this latency difference was apparent for both left utility (0.91s vs. 1.25s) and right utility (0.65s vs. 1.37s).

Therefore, in the preSMA, decisions and stimulus values are encoded by largely separate groups of neurons, with utility encoding appearing earlier than the decision. This time course and encoding scheme suggests that preSMA encodes pertinent stimulus features pre-decision, thereby revealing a potential substrate for value-based decision-making.

### Representation of decision conditioned variables

Representing the expected outcome of a choice is a critical step in decision making as it facilitates learning by way of comparison to observing the actual outcome received. We therefore next examine the neuronal representation of selected option’s utility (see full selection-based model in Table S2). Components of the selected option’s utility were encoded in vmPFC and dACC, but not in preSMA. In vmPFC, selected q-values, uncertainty, and novelty were encoded in both the trial onset and pre-decision period (Fig. 5 A-C, Fig. 5 E shows an example). In dACC, all three variables were also encoded in the pre-decision period. We also found that selected novelty neurons became active significantly earlier in vmPFC than dACC (−1.06s vs. dACC: −0.80s, p < 0.05), with no significant latency differences between the two areas for the other two variables (Fig. S4 A). Furthermore, we examined encoding of value for the rejected option (see Supplementary Material, Fig. S3 A-C). Additionally, a significant proportion of vmPFC selected uncertainty neurons signal whether a trial is exploratory or not prior to button press (see Supplementary Material). Overall, these findings indicate that single neurons in vmPFC and dACC encode value components contingent on the decision that was made.

**Figure 5:**
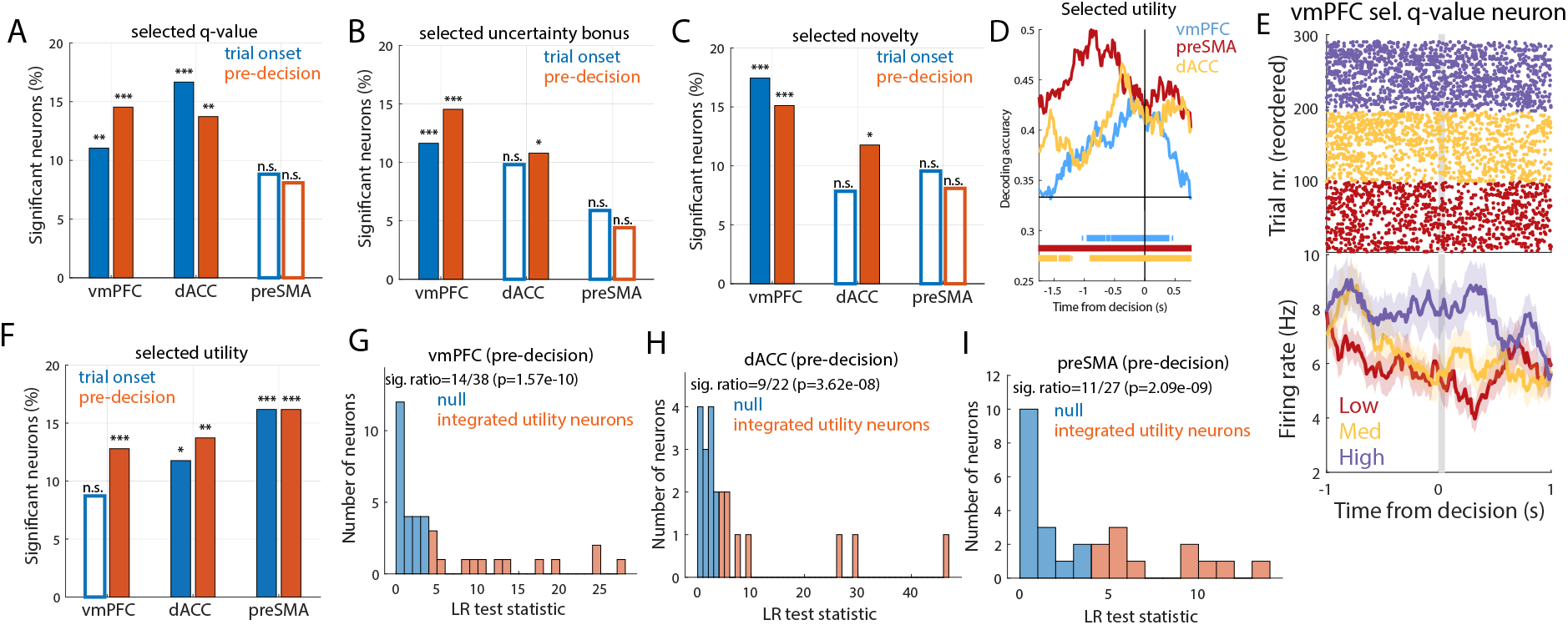
Encoding selected stimulus properties. (a) Percentage of selected q-value neurons in vmPFC, dACC, and preSMA, in the trial onset (blue) and pre-decision (orange) periods (* = p<0.05; ** = p<0.01; *** = p<0.001, binomial test). Hollow bars indicate non-sensitive counts. (b) Same, for selected uncertainty. (c) Same, for selected novelty. (d) dPCA selected utility decoding in the pre-decision period, for vmPFC (blue), preSMA (red), and dACC (yellow). Bars indicate significant decoding accuracies for each brain region, comparing to a bootstrapped null distribution. (e) VmPFC selected q-value neuron. Top: Raster plots. For plotting, we sorted trials by q-value tertiles (purple: high; yellow: medium; red: low). Bottom: PSTH (bin size = 0.2 s, step size = 0.0625 s). Gray bar indicates button press. Shaded areas indicate standard error. (f) Same as (a), for selected utility. (g) Histogram of likelihood ratio test statistics across candidate vmPFC integrated selected utility neurons (orange), in the pre-decision period. For the remaining neurons (blue) a null model containing only selected q-values was not rejected. (h) Same as (g), for dACC. (i) Same as (g), for preSMA.

Similar to the integrated positional utility analysis, we defined a group of candidate integrated selected utility neurons as the subset of units that correlated with the selected option’s q-value or utility (shown in Fig. 5 F). We determined that, in all brain areas, the number of neurons selected this way was larger than expected by chance (Figs. 5 G-I) (vmPFC: *p* < 10^−9^; dACC: *p* < 10^−7^; preSMA: *p* < 10^−8^). The activity of this subset of neurons was therefore indicative of an integrated selected utility.

Finally, we examined the points in time at which the selected option’s integrated utility could be decoded from pooled activity across all neurons in the regions of interest (using dPCA, see Materials and Methods). This analysis revealed robust decoding of selected utility in all brain areas, with a notably earlier onset in preSMA compared to both vmPFC and dACC (Fig. 5 D). Motivated by the earlier utility decoding in preSMA, we tested whether selected utility neuron latency times were also shorter in preSMA than in the other areas. A Poisson latency analysis revealed an onset time in preSMA of −0.83*s* ± 0.01, which was significantly earlier than in vmPFC (−0.79*s* ± 0.01, *p* < 0.05, one-sided rank sum test) and dACC (−0.71*s* ± 0.02, *p* < 10^−5^, one-sided rank sum test).

Taken together, these findings establish widespread value coding specific to the chosen option in all tested brain areas. One interpretation of these findings is that features of selected stimuli are monitored after the decision in the time window that immediately precedes the button press. While all areas displayed evidence of integrated selected utility coding, the preSMA represented this signal earlier than the other regions, consistent with the possibility that the preSMA is more closely involved in the choice process.

### Post-feedback neuronal responses

Behavioral consequences offer information that can be leveraged to make better decisions in the future. We tested for neurons encoding reward information, probing for representations of outcome, expected value and RPE during the feedback period. (Figs. S6 A-D). Outcome (win/lose) was robustly encoded in dACC, preSMA and vmPFC (Percentage of neurons selected 34.3%, *p* = 0; 35.3%, *p* = 0; and 17.4%, *p* < 10^−9^/3, respectively, binomial test). The q-value of the selected stimulus was encoded in vmPFC and preSMA, but not dACC (12.2%, *p* < 10^−4^/3 and 15.4%, *p* < 10^−5^/3, respectively).

We probed for two forms of the RPEs: the RPEs absolute value tracking surprise irrespective of valence, and a proxy for signed RPE, outcome minus selected q-value. Signed RPE was encoded in vmPFC and preSMA (11.6%, *p* < 10^−3^/3 and 16.9%, *p* < 10^−7^/3, respectively), but not dACC (Figs. S6 D). In contrast, we did not find significant numbers of neurons encoding the RPEs absolute value in either brain area (Figs. S6 C).

Latency analysis revealed that contrary to error signals we have studied previously,^38^ dACC neurons encoded outcome significantly earlier than both preSMA and vmPFC (Fig. S6 I; outcome-aligned median latency: 0.50s vs. 0.79s, *p* < 10^−17^ and vs. 0.81s, *p* < 10^−7^, two-sided rank-sum test). There was no difference between the onset of outcome signals in preSMA and vmPFC (*p* = 0.21, rank-sum test). Lastly, outcome neurons became active earlier than selected q-value neurons in all three regions (median times: vmPFC: 0.91s, *p* < 0.05; dACC: 0.69s, *p* < 0.001; preSMA: 0.97s, *p* < 10^−4^, rank-sum test), indicating that selected q-value representations were not persistently maintained from the choice period.

Whereas the majority of neurons encoded one of these three variables exclusively, approximately one fourth encoded multiple variables (Figs. S6 K; proportion of mixed neurons out of all sensitive neurons: vmPFC: 19%; dACC: 16%; preSMA: 25%). We cautiously speculate that this mixed selectivity could be accounted for by independent probabilities of a neuron representing a given variable. In preSMA, the proportion of neurons that signaled both outcome and selected q-value was higher than expected by independence (*p* < 0.05, Jaccard index test), but not in vmPFC (*p* = 0.07, Jaccard index test) or dACC (*p* = 0.18, Jaccard index test). This suggests that the activity of individual neurons in preSMA uniquely contains sufficient information to support the computation of reward prediction errors.

## Discussion

We investigated value-based decision making in human single neurons, while manipulating variables relevant to resolution of the explore/exploit dilemma; specifically, stimulus value, uncertainty and novelty. By recording from three areas of the prefrontal cortex implicated in decision-making across humans and other animals,^9, 11, 21, 25, 33, 39–43^ we identified how these variables are encoded, and addressed how they are integrated to inform decisions. Our findings highlight a particularly important role for human preSMA neurons in value-based decisions.

We found evidence for separate representations of the EV, uncertainty and novelty associated with options under consideration in human single neurons in both the preSMA and vmPFC, supporting the separable encoding of each of these decision variables across these areas. Crucially, we also found that a subset of EV coding neurons were better explained by an integrated utility signal, in which the option’s EV was combined with uncertainty and novelty. This signal was most robustly represented in the preSMA, where it was encoded both at the single neuron and population levels. These findings provide a proof of principle for the existence of an integrated utility signal in human prefrontal neurons.

We also identified a distinct population of preSMA neurons encoding the decision itself above and beyond stimulus utility, expanding on previous findings linking preSMA to volitional decision making.^44^ Thus, unlike dACC or vmPFC, the preSMA represented not only the key utility signal which informs choice, but also the behavioral product of the decision itself. These results for value-based decisions expand on previous work which reported choice signaling in categorization and memory tasks in preSMA and dACC.^45^ We found robust outcome tracking in dACC and preSMA, in consonance with previous findings in human dmPFC,^46^ and while preSMA (as well as SMA^47^) had been shown to monitor internally generated error responses, preceding dACC error neurons temporally,^38^ we observed that value-based feedback elicited earlier outcome responses in dACC than in preSMA. Reward prediction error, on the other hand, was more robustly encoded in preSMA than in dACC. Taken together, these results appear to position the preSMA as playing a central role in value-based decision-making in humans, particularly in decision tasks that elicit the integration of multiple stimulus features as is required to balance the explore/exploit trade-off. Although we found that the preSMA plays a privileged role in encoding decision variables, we expect that these computations are likely supported by a broader cortico-striatal network beyond the preSMA alone.^48–54^

Our findings support a distinction between dorsal and ventral areas of cortex, whereby dorsal regions contribute to action-based decisions while more ventral areas such as the vmPFC are involved in valuation but not in decisions over actions.^33, 40, 55–59^ Here we find that a similar organization applies at the level of human single neurons. However, we also found a degree of specificity within the dorsal human prefrontal cortex as to where integrated utility and the decision itself are encoded: in preSMA but not as robustly in dACC. These findings situate the human preSMA as being more prominently involved in the computations directly required for value-based decision-making than the sub-region of dACC from which we recorded. The present findings thus contribute to a more fine-grained understanding of functional specificity within dorsomedial prefrontal cortex.

We also looked for the representation of variables pertinent to the selected option; and thus, contingent on the decision made. The integrated utility for the option that was ultimately chosen was found to be widely encoded throughout all three of the brain regions we recorded. It is noteworthy that this signal emerged markedly earlier in the preSMA than in vmPFC, consistent with the possibility that preSMA is more proximal to the generation of the decision itself than is the vmPFC. Unlike the preSMA, single neuron activity in vmPFC also correlated with individual decision variables for the value, uncertainty and novelty of those stimuli that had been selected on a given trial. Furthermore, a significant portion of vmPFC neurons encoding selected uncertainty were also modulated by whether a decision was classified as exploratory or not (Fig. S5). When taken together, these findings suggest a role for vmPFC neurons in post-decisional monitoring of option features, especially in the context of exploratory decision making.

We found widespread outcome encoding across all three regions, in consonance with a vast literature implicating prefrontal cortex in signaling outcomes, in rodents,^60–63^ monkeys,^64–67^ and humans.^41, 68^ We further found significant evidence for concurrent encoding of outcomes and selected EVs in preSMA post-feedback, which together constitute the two main components of reward-prediction errors.^1, 69, 70^ These findings suggest that preSMA neurons can support learning of reward expectations.

In conclusion, our results situate the human preSMA as an important center for value-based decision-making, with a robust encoding of decision variables and, most crucially, an integrated utility signal at the single neuron level that can be leveraged to inform choice. While vmPFC neurons encoded pre-decision variables as well as post-decision variables contingent on choice, neither this region nor the dACC showed an equivalently robust encoding of pre-decision integrated utilities or the choice itself. These findings suggest that value-based decision-making during exploration depends on highly specialized computations performed in distinct areas of the prefrontal cortex. Furthermore, the existence of an integrated utility at the level of single neurons that could serve as the input to the choice process suggests that relevant decision variables such as EV, uncertainty and novelty are first integrated into a unified neuronal representation prior to being entered into a decision comparison, shedding light on how subjective utility-based choices are implemented in the human brain.

## Acknowledgements

We would like to thank the members of the O’Doherty and Rutishauser labs for discussions and feedback. We thank all subjects and their families for their participation, as well as nurses and medical staff for their work. This work was supported by National Institutes of Health Grants R01DA040011 and R01MH111425 (to J.P.O.), R01MH110831 (to U.R.), and P50MH094258 (to J.P.O. and U.R.).

## Author Contributions

T.G.A., J.C., U.R., and J.P.O. designed the study. T.G.A. performed the experiments. T.G.A. and J.C. analyzed the data. T.G.A., J.C., A.N.M. U.R., and J.P.O. wrote the paper. A.N.M. performed surgery and supervised clinical work.

## Competing interests

None

## Data statement

Data and code will be deposited in a public archive upon acceptance of the manuscript.

## Notes

### Competing Interest Statement

The authors have declared no competing interest.

