## Supplemental Text for "Neurons in human pre-supplementary motor area encode key computations for value-based choice"

#### Supplementary Text

##### Model comparison

We compared how well two different computational models explained the observed behavior. Both models allowed for uncertainty and novelty to contribute to decision making beyond what could be implemented with a simpler reinforcement learning framework and also allowed incorporating patients' individual preferences. The compared models had two distinct mechanisms for how novelty, uncertainty, and q-value interacted to create stimulus utility (see Materials and Methods for detailed model descriptions). The first model we tested is a familiarity modulated upper confidence bound (fmUCB) model, shown to explain behavior from a neurotypical population in this task well[1] (Fig. 1 G). In this model, stimulus utility is equal to a linear combination of the stimulus q-value and an uncertainty bonus, defined as a weighted product between uncertainty and novelty, allowing for the uncertainty and novelty factors to interact. The second model we tested was a 'linear novelty model', in which stimulus utility is equal to a linear combination of q-value, an uncertainty bonus derived from stimulus uncertainty alone, and a novelty bonus derived from stimulus novelty alone, without direct interactions between novelty and uncertainty.

We performed model fitting and comparisons for the two models across the patient population using hierarchical Bayesian inference[2] (see Fig. S1 for values of fit parameters). The fmUCB model explained the observed behavior across the patient population significantly better, with an estimated model frequency of 78.3% and exceedance probability of 99.6%. This is consistent with the results of a study from a larger cohort of healthy participants performing the same task,[1] suggesting a non-linear interaction between uncertainty and novelty can drive decisions.

To test whether patient behavior reflected the instruction to reset the q-values and uncertainty bonuses at the beginning of every block as instructed, we also compared the fmUCB model with a 'no reset' model. This model was similar to the fmUCB model except that it did not reset q-value and uncertainty estimates from one block to the next. The fmUCB model had an exceedance probability of 99.9% compared to the no-reset model, indicating that patients reset contingencies between blocks as instructed. Lastly, we also compared the fmUCB model with a simpler reinforcement learning model in how well they could recover decision behavior. This analysis showed that a simple RL model did not adequately capture the observed behavioral effects beyond seeking expected value.

These behavioral modeling results show that a fmUCB mechanism is appropriate to model the behavior of our subjects. For all subsequent neural data analysis we therefore used the variables derived from the fmUCB model as regressors.

#### Additional behavioral analysis

For the chosen fmUCB model, we summarized its parameter fits to gain more insight into the behavior of the patients as a group (Fig. S1 A). The softmax inverse temperature was $4.6 \pm 0.47$ and the learning rate was $0.36 \pm 0.03$. The novelty intercept parameter represents a novelty initiation (nI) bias and was negative ($-0.30 \pm 0.13$, $p < 0.05$, t-test), indicating a slight preference for familiar stimuli in the early trials of a block. The uncertainty intercept (uI) parameter represents how much value was assigned to uncertainty in the beginning of a block. This parameter was negative ($-0.10 \pm 0.01$, $p < 10^{-5}$, t-test), indicating a slight uncertainty avoidance early in the block. The uncertainty terminal value parameter, which represents the value assigned to uncertainty at the end of a block, was also negative ($-0.15 \pm 0.03$, $p < 10^{-4}$, t-test), indicating that uncertainty was still valued negatively on average by the end of the blocks.

To ensure that the fmUCB model correctly reproduces the effects of expected value, uncertainty and novelty observed in our subjects, we performed a posterior predictive check analysis (Fig. S1 B-C). We exposed the fmUCB model to the same sequence of trials each patient experienced, with each patient's model fits, and generated a decision for each trial used the decision probabilities inferred from the model. We then used these choices to fit a logistic regression which maps the effects of the various variables on the choices (equivalent to Fig. 1 F). To account for variability in probabilistic decisions, we repeated this procedure 100 times and generated a distribution of regression coefficient estimates, which we compared to the actual effects observed in the subjects. For comparison, we performed the same procedure with an equivalent model (a simple reinforcement learning model), except that it did not model any effects of uncertainty and novelty, including only the softmax beta and learning rate parameters. To quantify the difference in how these models recovered effects of expected value, uncertainty and novelty on behavior, we computed the 95% confidence intervals for the logistic regression estimates of value feature effects on actual patient decisions (as displayed in Fig. 1, see Materials and Methods). Then, we obtained the proportion of overlap between these confidence intervals and the recovered effect estimates obtained from 100 iterations of simulated sessions in the posterior predictive check analysis. Using the simple RL model, we observed an overlap of $[95\%, 50\%, 14\%]$ for the estimates of expected, uncertainty and novelty, respectively. With the fmUCB model, we obtained overlaps of $[89\%, 88\%, 85\%]$. Therefore, the fmUCB model captured the behavioral effects of uncertainty and novelty better, while a simpler reinforcement learning model was not appropriate to capture the observed behavioral effects beyond seeking expected value.

#### Encoding rejected values

Models of decision making often also depend on maintaining information about the value of options not chosen to evaluate the outcome received. We therefore also examined whether rejected q-values, uncertainty bonuses and novelties were represented (Fig. S3 A-C). Unlike selected q-values, preSMA did represent rejected q-values (trial onset 14.0%, $p < 10^{-4}/6$:, pre-decision: 16.9%, $p < 10^{-6}/6$, binomial test, Bonferroni corrected). Novelty of the rejected option was encoded in vmPFC (trial onset 11.6%, $p < 0.001/6$:, pre-decision: 16.8%, $p < 10^{-7}/6$, binomial test, Bonferroni corrected).

#### Exploratory signaling in vmPFC uncertainty neurons

One potential reason for representing the uncertainty of the selected option is to enable exploratory decision making, which would entail deliberately choosing an item with lower q-value. We therefore divided trials into putative explore and non-explore categories. Trials in which the patient chose the option which had the lower q-value but the higher uncertainty bonus were classified as putative explore trials, while all others were classified as non-explore trials (Fig. S5 A).

Then, in the sub-populations of vmPFC and dACC neurons in the pre-decision period that were sensitive to selected uncertainty, we performed a Poisson GLM analysis using the explore trial flag as a regressor, correcting for selected uncertainty as a regressor of no interest. We subsequently tested whether neurons whose activity was significantly modulated by the explore flag significantly overlapped with the sub-populations of vmPFC and dACC neurons that encoded selected uncertainty (Fig. S5, B). We found a significant overlap in vmPFC ($p < 0.01$, Jaccard index test), but not in dACC ($p = 0.13$, Jaccard index test). Therefore, a significant proportion of vmPFC selected uncertainty neurons signal whether a trial is exploratory or not prior to button press.

### Materials and Methods

#### Electrophysiology and recording

We used Behnke-Fried hybrid depth electrodes (AdTech Medical), positioned exclusively according to clinical criteria (Supplementary Table S3). Broadband extracellular recordings were performed with a sampling rate of 32 kHz and a bandpass of 0.1-9000Hz (ATLAS system, Neuralynx Inc.). The data set reported here was obtained bilaterally from ventromedial prefrontal cortex (vmPFC), dorsal anterior cingulate cortex (dACC), and pre-supplementary motor area (preSMA) with one macroelectrode on each side. Each macroelectrode contained eight 40 $\mu$m microelectrodes. Recordings were bipolar, utilizing one microelectrode in each bundle of eight microelectrodes as a local reference.

#### Patients

Twenty patients (fourteen females) were implanted with depth electrodes for seizure monitoring prior to potential surgery for treatment of drug resistant epilepsy. Two of the patients performed the task twice, totalling 22 recorded sessions. Human research experimental protocols were approved by the Institutional Review Boards of the California Institute of Technology and the Cedars-Sinai Medical Center. Electrode location was determined based on preoperative and postoperative T1 scans obtained for each patient.

#### Task

Patients performed a two-armed bandit task (Fig. 1 B). The task contained 20 blocks of 15 trials, for a total of 300 trials. The 20 blocks were split into 2 recording sessions with 10 blocks each, with a 5 minute break in between sessions. Each trial began with a baseline period (sampled randomly from a uniform distribution of 0.75-1.25s), followed by a choice screen showing the two available slot machines presented on the left or on the right of the screen. The identity of each slot machine was uniquely identifiable by a painting displayed on the center of each slot machine. Patients had to decide between the left or the right option by pressing a button in less than 3s, or the trial would be considered missed and no reward would be accrued. Across all trials, mean reaction time (RT) was $1.47s \pm 0.02$ (relative to onset of choice screen). Following the button press, the chosen slot machine is shown for a period of 1-2s (sampled randomly from a uniform distribution), followed by the outcome screen shown for 2s. The outcome screen showed either a golden coin to represent winning a reward, or a crossed out coin to represent not winning (both shown on top of the chosen slot machine).

To shape the novelty and uncertainty of presented stimuli, we manipulated which stimuli would appear in each block and each trial according to the rules described as follows. For each block, the identity of the two slot machines that appeared in each trial was drawn randomly from a set of 3 possible options, selected specifically for each block. In the first block, the 3 options were selected randomly from a set of 200 paintings. In every subsequent block, one out of the three stimuli from the previous block was chosen to be replaced, substituting it for a novel unused stimulus out of the 200 paintings.

To manipulate the interaction between stimulus novelty and trial horizon, in every block after the first one, we chose stimuli to be held out and only presented after a minimum trial threshold, selected randomly for each block, between 7 and 15 trials. For every block after the first one, we alternated whether the held out stimulus would be one of the familiar ones or the novel stimulus for that block.

The probabilities of receiving a reward from each slot machine were reset in the beginning of every block, and determined according to the chosen difficulty of each block, which alternated between the easy and hard conditions. Crucially, these reward probabilities did not change within each block. In the easy condition, reward probabilities were more widely spaced out between different slot machines, and chosen from the values $[0.2, 0.5, 0.8]$. In the hard condition, the possible probabilities were $[0.2, 0.5, 0.6]$.

Some patients performed a shorter variant of the task, which consisted of 206 trials across 10 blocks (see Supplementary Table S1). In this version, the set of possible stimuli in each block contained 5 options, in each block after the first one 2 novel options were introduced, one of which composed the held out set along with one out of the 3 familiar options from the previous blocks. Bandit win probabilities were sampled from the linearly spaced interval $[0.2, 0.8]$ in easy blocks and from $[0.4, 0.6]$ in hard blocks. We pooled data from the two task variants together for all analysis.

#### Behavioral analysis and computational modeling

##### Logistic regression for value components and decisions

We used a logistic regression model to describe how the past history of rewards, sampling history, stimulus exposure history, and their interactions with trial number correlated with decisions (Fig. 1 F). For this, we defined q-value $Q_s$ as the mean of a beta distribution which estimates the probability of receiving a reward from a bandit, as determined by the history of wins and losses after sampling a stimulus $s$, as well as $\delta Q = Q_{left} - Q_{right}$, the difference between left and right Q-values. Similarly, we define an uncertainty value $U$ as the variance of the same beta distribution, as well as its corresponding differential $\delta U = U_{left} - U_{right}$. Finally, we defined novelty $N$ as the variance of a beta distribution in which $\beta = 1$ and the $\alpha$ parameter is the number of times patients were exposed to a stimulus $s$ in the entire session, as well as its corresponding differential $\delta N = N_{left} - N_{right}$:

We then performed a logistic regression using MATLAB's function *mnrfit* to model the probability $p_{left}$ of a left decision based on these regressors as well as their interaction with the trial number $t$

within a block:

$$\log \frac{p_{left}}{1 - p_{left}} = \beta_0 + \beta_1 \delta Q + \beta_2 \delta U + \beta_3 \delta N + \beta_4 \delta Q \cdot t + \beta_5 \delta U \cdot t + \beta_6 \delta N \cdot t \tag{1}$$

**Familiarity gating model of exploration (fmUCB)**

We compared two computational models fit to patients' behavior. Individualized model fits and model comparisons were obtained across the patient population through hierarchical Bayesian inference.[2] This method yielded model parameters for each subject in the data set, for each of the tested models, as well as exceedance probabilities, which expressed the probability that either model was the most frequent in the behavioral dataset.[3]

The first model we tested is a fmUCB model[1] of exploratory decision making. In this model, the choice probability for a decision $d$ in a trial $t$ is estimated using the utilities assigned to the left ($U_L$) and right ($U_R$) options, through a softmax function:

$$p_t(d = LEFT) = \frac{1}{1 + e^{\beta(U_{R,t} - U_{L,t})}} \tag{2}$$

In this equation, $\beta$ is the inverse temperature free parameter. To balance incentives to explore and exploit different stimuli, the utilities assigned to each stimulus $s$ on a trial $t$ were defined to be the sum of its weighted q-values and an uncertainty bonus $B$, depending on the past history of rewards received from the slot machine, and how often the slot machine had been sampled, respectively:

$$U_{s,t} = Q_{s,t} + B_{s,t} \tag{3}$$

The q-value was defined similarly to the expected value of a beta distribution, as a function of the past history of wins and losses received from a slot machine, modified to account for the effect of recency over stimulus preferences:

$$Q_{s,t} = \frac{\alpha_{s,t}}{\alpha_{s,t} + \beta_{s,t}} \tag{4}$$

In this equation, $\alpha_{s,t}$ and $\beta_{s,t}$ describe the effect of previous wins and previous losses, respectively, received from the slot machine $s$ before trial $t$.

The $\alpha$ term is defined as follows, where $H^W_{s,t}$ is how many times sampling slot machine $s$ has resulted in a win before trial $t$, and $w$ is an exponentially decaying effect of recency. The time scale of this exponential decay is determined by a learning rate free parameter $\lambda$, fit in the interval (0,1):

$$\alpha_{s,t} = 1 + \sum_{i=1}^{t-1} w_{i,t} H^W_{s,t} \tag{5}$$

$$w_{i,t} = (1 - \lambda)^{(t-i)} \tag{6}$$

Similarly, the $\beta$ term is defined as follows, where $H^L_{s,t}$ is how many times sampling slot machine $s$ has resulted in a no-win before trial $t$:

$$\beta_{s,t} = 1 + \sum_{i=1}^{t-1} w_{i,t} H^L_{s,t} \tag{7}$$

We also allowed novelty to bias the initialization of the $\alpha$ and $\beta$ hyperparameters, to include an optimistic initialization strategy[4] for exploration. This was done by including a novelty initialization bias free parameter $n_I$, which was modulated by the same exponential decay $w_{0,t}$, creating the novelty bias $n_I w_{0,t}$. If $n_I w_{0,t} > 0$, we would add this quantity to $\alpha_{s,t}$, resulting in a novelty seeking bias, and if $n_I w_{0,t} < 0$, we added this quantity to $\beta_{s,t}$, resulting in a novelty avoidance bias.

The uncertainty bonus term in Equation 3 was defined as a function of raw stimulus uncertainty, gated by familiarity, and weighed by each patients' uncertainty preferences, according to the weight parameter $w^U_t$, as a function of the trial horizon within a block, as will be defined further:

$$B_{s,t} = V_{s,t} F_{s,t} w^U_t \tag{8}$$

Raw stimulus uncertainty $V_{s,t}$ was defined similarly to the variance of a beta distribution, as a function of how many times a stimulus has been sampled, using the previously defined $\alpha_{s,t}$ and $\beta_{s,t}$ terms:

$$V_{s,t} = 12 \frac{\alpha_{s,t}\beta_{s,t}}{(\alpha_{s,t} + \beta_{s,t})^2(\alpha_{s,t} + \beta_{s,t} + 1)} \tag{9}$$

We introduced a normalizing factor of 12 to the raw stimulus uncertainty equation to ensure that maximal uncertainty, obtained when $\alpha_{s,t} + \beta_{s,t} = 1$, is equal to 1.

The familiarity gating was introduced to allow for the novelty of a stimulus, as a function of how many times it has been seen throughout the session, to interact with the behavioral effects of uncertainty. Defining $g$ as a familiarity gating free parameter, fit for each subject, the familiarity gating $F_{s,t}$ is defined as follows:

$$F_{s,t} = 1 - g N_{s,t} \tag{10}$$

In this equation, $N_{s,t}$ is the novelty value for slot machine $s$ in trial $t$, defined as a monotonically decreasing function of the number of exposures for $s$, defined as $E_{s,t}$:

$$N_{s,t} = 12 \frac{E_{s,t} + 1}{(E_{s,t} + 2)^2(E_{s,t} + 3)} \tag{11}$$

This definition of novelty was chosen to create a similar functional form to the uncertainty value, while enforcing that maximal novelty, for stimuli that had not been exposed before, was equal to 1.

Finally, to allow for switching between exploration and exploitation within a block, the effect of trial horizon over the uncertainty bonus was defined a linear function of the trial number within a block, adding the free parameters for terminal uncertainty $uT$ and uncertainty intercept $uI$, where $T$ is the maximal trial horizon for a block and $uS$ is the uncertainty slope:

$$uS = \frac{(uT - uI)}{T} \tag{12}$$

$$w_t^U = uI + uS(t-1) \tag{13}$$

**Alternative model with independent utility of novelty (linear novelty model)**

In the second model we tested, uncertainty and novelty did not interact directly in the construction of the uncertainty bonus, but are added as independent components of the stimulus utility value. Concretely, Equation 3 is modified to add a novelty bonus $N^*_{s,t}$ to utility:

$$U_{s,t} = Q_{s,t} + B^*_{s,t} + N^*_{s,t} \tag{14}$$

The uncertainty bonus from Equation 8 was adapted, creating a modified uncertainty bonus $B^*_{s,t}$, to remove the interaction with novelty through familiarity gating:

$$B^*_{s,t} = V_{s,t} w_t^U \tag{15}$$

We defined the novelty bonus $N^*_{s,t}$ similarly to the uncertainty bonus, by multiplying the previously defined novelty value (Equation 11) by a novelty weight free parameter ($w_t^N$), which was fit for each patient:

$$N^*_{s,t} = N_{s,t} w_t^N \tag{16}$$

The remaining components of the alternative model with a novelty bonus are kept the same as in the fmUCB model.

#### Neural data pre-processing

We performed spike detection and sorting with the semiautomatic template-matching algorithm OSort.[5] Channels with interictal epileptic activity were excluded. Across all 22 sessions, we obtained 191 vmPFC, 137 preSMA and 108 dACC putative single units (436 total). In this manuscript we refer to these isolated putative single units as "neuron" and "cell" interchangeably. For the single neuron encoding analyses in this study we pre-selected only neurons with more than 0.5Hz average firing rate across all trials, resulting in 172 vmPFC, 136 preSMA and 102 dACC putative single units (410 total).

#### Poisson GLM encoding analysis

We used Poisson regression GLMs to select for neurons, with response variable the number of spikes fired and the dependent variable different subsets of model variables. We computed the spike counts in every trial in four windows of interest (trial onset, from 0.25s to 1.75s, aligned to trial onset; pre-decision, from -1s to 0s, aligned to button press; and outcome, from 0.25 to 1.75s, aligned to outcome onset). For visualization purposes, we also fit the same models with 0.5s time windows, sliding by 16ms steps, within the same time limits. We then tested hypotheses about how the spike count of each neuron was correlated with left and right utility ($U_L$,$U_R$), chosen side ($Side$), left and right q-value ($Q_L$,$Q_R$), left and right uncertainty bonus ($B_L$,$B_R$), left and right novelty ($N_L$,$N_R$), as well as their selected and rejected counterparts, outcome ($O$) and absolute reward prediction error ($|RPE|$). Additionally, we performed these encoding analyses utilizing the raw uncertainty values $V_{s,t}$ instead of the transformed uncertainty bonus values $B_{s,t}$, and obtained equivalent results.

We also tested whether neuronal activity in the pre-decision period correlated with whether a trial was classified as an explore or a non-explore trial, correcting for selected uncertainty bonus. We defined explore trials as those in which $Q_{sel} < Q_{rej}$ and $U_{sel} > U_{rej}$, defining the explore flag $Explore = 1$ for those trials and $Explore = 0$ for all others. For these analyses, we specified the models described on Table S2 and fit them with the MATLAB function $fitglm$.

To understand the overall role of q-values, uncertainty and novelty regardless of the position of stimuli on the screen (Fig. 2), we fit the full positional model coefficients and reported the proportion of sensitive neurons for the left and right components together, as positional q-value neurons, positional uncertainty bonus neurons and positional novelty neurons.

For the outcome analysis (Fig. S6), we performed an F-test for the difference between coefficients for outcome and selected q-value ($b_1 - b_2$), as a proxy for reward prediction error coding, and reported the proportion of neurons for which the contrast is different than 0. We also report the number of neurons whose activity correlates with outcomes and absolute reward prediction error at the time of outcome.

#### Poisson latency analysis

To determine when individual neurons became active at a single trial level, we performed Poisson latency analyses[6] for pre-selected groups of neurons sensitive to the variable of interest in the encoding analyses (Figs. 2 G-H; Fig. 5 B; Fig. S6 H). This method detects the first point in time in which interspike intervals significantly differ from what would be expected from a constant firing rate Poisson point process, using the neuron's average firing rate as the rate parameter. We used a significance parameter of $p < 0.05$ as our burst detection threshold for all analyses.

#### Jaccard index test

After performing Poisson GLM encoding analyses, we tested whether the sub-populations of neurons which were sensitive to two variables of interest had significant overlap. For this, we computed the Jaccard index[7] of overlap between neurons sensitive to each of the variables $X$ and $Y$, where $N_X$ and $N_Y$ indicate the number of neurons sensitive to the variables X and Y, respectively, and $N_{X,Y}$ indicates the number of neurons concurrently sensitive to both variables:

$$J = \frac{N_{X,Y}}{N_X + N_Y - N_{X,Y}} \tag{17}$$

To compute p-values for each comparison between two variables, we bootstrapped a null distribution of Jaccard indexes using 1000 reshuffles, considering X and Y are independent variables with a false positive rate of $p = 0.05$.

#### Likelihood ratio hypothesis testing

We tested whether neurons in positional q-value or positional utility sensitive sub-populations had their activity better explained by an unrestricted model including the main additive components of utility (q-value and uncertainty bonus) or by a restricted model including only q-values, given the correlations we observed between q-values and integrated utility values. Neurons which had their activity better explained by the unrestricted model were defined as true integrated utility neurons.

Before constructing the unrestricted and restricted models, we determined the preferred side of each neuron by fitting their activity with the utility and decision model, including left utility, right utility and decision as regressors (Table S2) and defining the preferred side as the one in which its utility regressor has the highest absolute t-score.

Then, using the spike count $Y$ of each neuron we fit an unrestricted GLM including q-values and uncertainty bonuses. We performed the model fitting and obtained a log-likelihood $L_u$ using MATLAB's function $fitglm$:

$$\log(E(Y|x)) = b_0 + b_1 Q_{preferred} + b_2 B_{preferred} + b_3 Decision \tag{18}$$

To each neuron in this sub-population we also fit a restricted GLM including q-values but not uncertainty bonuses and obtained its log-likelihood $L_r$:

$$\log(E(Y|x)) = b_0 + b_1 Q_{preferred} + b_2 Decision \tag{19}$$

Finally, we performed likelihood ratio tests, with MATLAB's function *lratiotest*, between the unrestricted and restricted models, by computing the likelihood ratio test statistic $LR = 2(L_u - L_r)$, and comparing it to a chi-squared null distribution for LR with one degree of freedom, stemming from one variable restriction. Neurons that rejected the null restricted model at a significance level of $\alpha = 0.05$ were defined as integrated utility neurons.

For the sub-population of integrated utility neurons, we used their fits from the unrestricted models to determine whether activity in these neurons correlated with q-values and uncertainty bonuses individually more than expected by chance. We averaged absolute t-scores for q-value and uncertainty bonus across integrated utility neurons to measure their collective degree of correlation regardless of excitation or inhibition. We then compared these values with average absolute t-scores obtained from bootstrapping 500 iterations of unrestricted model fits shuffling spike counts $Y$. We derived p-values from the number of times the true average absolute t-score surpassed the bootstrapped iterations.

Similarly, we performed a likelihood ratio test to test whether neurons encoded an integrated selected utility signal in the pre-decision period by fitting the following unrestricted model:

$$\log(E(Y|x)) = b_0 + b_1 Q_{selected} + b_2 B_{selected} \tag{20}$$

Subsequently, we compared the unrestricted model with the following null restricted model:

$$\log(E(Y|x)) = b_0 + b_1 Q_{selected} \tag{21}$$

We then followed the same likelihood test protocol described above to determine whether neurons would be classified as integrated utility neurons or not.

#### Dimensionality reduction and decoding with dPCA

To decompose the contribution of variables of interest and decisions to the neural population data and decode these variables interest from patterns of neural activity, we employed demixed principal component analyses (dPCA).[8]

For each variable of interest, and each brain area, we created a pseudopopulation aggregating trials from all patients in order to generate a full data matrix $X$, with dimensions $(N, SQTK)$, where N is the total number of neurons recorded in that brain area, S is the number of stimuli quantiles used to partition trials (3: low, medium, and high), Q is the number of possible decisions (2: left and right),

T is the number of time bins, and K is the number of trials used to construct the pseudopopulation as described further. Firstly, we binned spike counts into $500ms$ bins, with a $16ms$ time window step. We repeated the binning procedure in two different time periods: the trial onset period (0s,2s), aligned to trial onset; and the pre-decision period (-2s,1s), aligned to button press.

#### Constructing pseudopopulations

To create neural pseudopopulations for dPCA, we pooled trials from all sessions and treated them as if they had been recorded simultaneously. To allow for trials from different sessions to be grouped together, despite having continuous variables of interest, we pooled groups of trials into 3 quantiles with the same number of trials, dividing the full range of each variable for each session into low, medium and high levels. After obtaining these quantiles, we assigned every trial in each session to one out of $3 \cdot 2 = 6$ categories, to account for all possible combination of quantile levels and decisions, and randomly sampled an equal number $k$ of trials from each category, for each session, such that $\sum_{sessions} k = K$. We chose $k = 15$, for it to be small enough to allow sampling an equal number of trials from each of the 6 categories for every session, while including as many training examples as possible.

To mitigate biases introduced during the random trial sampling procedure, we repeated these steps 10 times, yielding 10 pseudopopulations, on which the dimensionality reduction and decoding procedures were repeated independently.

#### dPCA dimensionality reduction

For dPCA dimensionality reduction, the full data matrix $X$ is centered over each neuron and decomposed as a factorial ANOVA, where $t$, $s$, and $d$ are labels to indicate the time, stimulus and decision marginalizations, respectively:

$$X = X_t + X_{ts} + X_{td} + X_{tsd} + X_{noise} = \sum_{\phi} X_{\phi} + X_{noise} \tag{22}$$

The goal of dPCA is then to minimize the regularized loss function, where F indicates the Frobenius norm and $\mu$ is the ridge regression regularization parameter, determined optimally through cross-validation:

$$L = \sum_{\phi} (||X_{\phi} - F_{\phi} D_{\phi} X||_F^2 + \mu ||F_{\phi} D_{\phi}||_F^2) \tag{23}$$

$F_{\phi}$ and $D_{\phi}$ are the non-orthogonal encoder and decoder matrices, respectively, arbitrarily chosen to have 3 components for each marginalization. Therefore, dPCA aims to reduce the distance between each marginalized data set and their reconstructed version obtained by projecting the full data matrix onto a low-dimensional space with the decoders D and reconstructing it with the encoders F.

#### dPCA decoding

We used the same dPCA framework to perform population decoding of the variables of interest. The dPCA linear decoding pipeline has been previously described in detail,[8] but we will briefly discuss it here.

Firstly, the pseudopopulation data matrix X of dimensions $(N, SQTK)$ is divided into train and test datasets by leaving out one random trial for each of the $SQ$ possible combinations of stimulus levels and chosen side, for all neurons and time points, to form $X_{test}$ of dimensions $(N, SQT)$ and $X_{train}$ with the remaining data points. We perform this random trial sampling procedure 100 times for each of the 10 random pseudopopulations, resulting in a total of 1000 random resamples.

We performed the aforementioned dPCA steps with the train data matrix $X_{train}$ to obtain a decoder matrix $D_{\phi,i}$, with $i = (1, 2, 3)$ representing each of the three demixed principal components for each marginalization $\phi$.

To perform stimulus decoding, we iterate over the three components $i = (1, 2, 3)$, to obtain the mean projections over all train trials, for each stimulus class $s = (1, ..., S)$ pertaining to the current marginalization, and the vectors of decoded projections for test trials, for each unique test trial $k = 1, ..., SQ$, representing all the possible stimulus-decision combinations:

$$P_{\phi,s}^{train} = \begin{bmatrix} < D_{\phi,1} X_{train} >_s \\ < D_{\phi,2} X_{train} >_s \\ < D_{\phi,3} X_{train} >_s \end{bmatrix}, P_{\phi}^{test,k} = \begin{bmatrix} D_{\phi,1} X_{test,k} \\ D_{\phi,2} X_{test,k} \\ D_{\phi,3} X_{test,k} \end{bmatrix} \tag{24}$$

We then defined the decoded class $C_k$ to be the one which minimizes the three-dimensional Euclidean distance between the test projection and the mean train projections:

$$C_k = \arg\min_s ||P_{\phi,s}^{train} - P_{\phi}^{test,k}|| \tag{25}$$

We obtained classification accuracy values for each trial resample by counting how many test trials were correctly labeled, and averaged classification accuracy values over the 100 random test trial resamples, as well as the 10 pseudopopulation resamples.

Equivalently, to perform decision decoding, we follow the same steps, except that we obtain mean projections over all train trials for each decision class $q = (1, ..., Q)$ to compare with test trial projections.

Significance scores for each time bin were determined by obtaining the distribution of null scores from the random test trial reshuffles, and computing the quantile placement of the true decoding accuracy, assuming an approximate normal distribution for reshuffled decoding accuracies. We subsequently Bonferroni corrected significance scores for multiple comparisons across time bins.

##### dPCA component projection distance

To summarize how dPCA representations of utility and decision differ for low/high utility trials, as well as left/right decisions (Fig. 4 F), for every time bin, we projected data $X_{subset}$ from each trial subset (low utility trials, high utility trials, left decision trials, and right decision trials) onto the demixed principal components, expressed by the decoder matrix D, obtaining $DX_{subset}$. Note that each row of D represents one demixed principal component for the dataset. We then computed Euclidean distances between projections $D_{decision} = ||DX_{left} - DX_{right}||^2$, and $D_{utility} = ||DX_{high} - DX_{low}||^2$. We subsequently normalized projection distances into the [0,1] range.

### Supplementary Tables

| Task version | Patients who performed it |
|---|---|
| Longer (300 trials) | P60,P61,P62,P63,P64,P65,P67,P69,P70,P71 |
| Shorter (206 trials) | P41,P43,P48,P49,P51,P54,P55,P56 |

Table S1: Patients who performed the longer (300 trials) or shorter (206 trials) version of the task. For all behavioral and neural analyses, datasets from both task versions were pooled.

| **Name** | **Model** | **Periods** |
|---|---|---|
| Full positional model | $\log(E(Y\|x)) = b_0 + b_1 Q_L + b_2 Q_R + b_3 B_L + b_4 B_R + b_5 N_L + b_6 N_R$ | trial onset, pre-decision |
| Full selection-based model | $\log(E(Y\|x)) = b_0 + b_1 Q_{sel} + b_2 Q_{rej} + b_3 B_{sel} + b_4 B_{rej} + b_5 N_{sel} + b_6 N_{rej}$ | trial onset, pre-decision |
| Selection-based utility model | $\log(E(Y\|x)) = b_0 + b_1 U_{sel} + b_2 U_{rej}$ | trial onset, pre-decision |
| Explore flag model | $\log(E(Y\|x)) = b_0 + b_1 Explore + b_2 U_{sel}$ | pre-decision |
| Decision and utility model | $\log(E(Y\|x)) = b_0 + b_1 U_L + b_2 U_R + b_3 Decision$ | trial onset, pre-decision |
| Outcome model | $\log(E(Y\|x)) = b_0 + b_1 O + b_2 Q_{sel} + b_3 \|RPE\|$ | outcome |

Table S2: Models for Poisson GLM single neuron encoding analysis.

| Patient ID | Left dACC | Right dACC | Left preSMA | Right preSMA | Left vmPFC | Right vmPFC |
|---|---|---|---|---|---|---|
| P71CS | -1.00,27.97,27.02 | 7.68,28.87,23.68 | -1.89,13.51,38.03 | 5.09,11.30,39.61 | -3.04,32.45,-0.64 | 5.06,34.29,6.30 |
| P70CS | -0.84,23.06,26.04 | 4.25,22.06,22.63 | -9.45,11.24,46.67 | 12.40,11.55,46.60 | -0.20,26.30,-6.79 | 10.50,29.65,-3.63 |
| P69CS | -6.13,27.44,22.69 | -1.66,27.37,22.25 | -2.02,23.89,39.02 | 2.92,32.34,40.50 | -3.73,38.30,-11.69 | 2.26,29.24,-9.80 |
| P67CS | -4.82,27.63,-11.69 | -0.27,28.18,-9.13 | -2.63,10.61,37.48 | 7.38,11.36,36.23 | -4.70,27.32,-14.06 | -1.21,28.12,-7.68 |
| P65CS | -4.66, 24.50, 24.48 | 5.66, 30.60, 25.28 | -0.82, 10.87, 52.78 | 1.53, 18.10, 48.52 | -3.69, 45.06, -12.11 | 6.46, 30.94, -11.70 |
| P64CS | -7.45, 35.50, 22.73 | 5.54, 32.54, 30.78 | -6.19, 21.34, 47.67 | 7.46, 22.79, 44.56 | -8.18, 30.42, -13.39 | 6.21, 31.14, -12.75 |
| P63CS | -5.57, 27.44, 26.20 | 11.66, 34.65, 33.43 | -8.53, 10.02, 41.57 | 9.46, 28.48, 48.52 | -3.52, 45.04, -14.44 | 9.54, 37.31, -13.49 |
| P62CS | NaN | 9.02, 15.72, 29.33 | NaN | 5.69, 16.28, 49.18 | NaN | 11.02, 34.13, -10.05 |
| P61CS | 0.00, 19.15, 22.74 | 6.82, 20.21, 27.63 | -4.55, 14.50, 38.70 | 14.92, 12.05, 36.59 | -3.40,41.84,-15.66 | 6.99,31.03,-12.39 |
| P60CS | -8.66, 30.23, 27.13 | 5.62, 31.09, 20.05 | -8.70, 19.60, 44.90 | 5.55, 26.23, 42.00 | -1.49,31.63,-16.97 | 2.48,33.28,-20.24 |
| P56CS | -7.01, 23.11, 30.17 | 16.27, 24.79, 25.90 | -6.92, 6.43, 46.70 | 29.08, 13.01, 45.88 | -6.96,34.62,-13.05 | 27.02, 31.69, -5.48 |
| P55CS | -4.14, 28.97, 24.60 | 3.20, 28.05, 21.05 | -3.88, 11.08, 46.18 | 7.58, 13.24, 42.29 | -8.64, 36.74, -6.98 | 6.32, 35.04, -11.70 |
| P54CS | -3.85, 29.02, 23.70 | 7.93, 33.96, 26.74 | -5.98, 20.87, 49.08 | 10.72, 26.92, 48.06 | -3.53, 36.85, -16.95 | 3.94, 31.75, -14.51 |
| P53CS | 0.95, 32.02, 27.76 | 9.47, 36.81, 26.07 | -4.30, 12.32, 51.06 | 10.21, 18.90, 49.06 | -0.97, 35.20, -17.12 | 6.83, 31.74, -14.96 |
| P51CS | -5.48, 31.63, 27.98 | 6.65, 30.73, 25.95 | -8.81, 8.15, 44.07 | 11.91, 9.98, 47.01 | -1.03, 36.31, -10.91 | 4.19,30.23,-15.23 |
| P49CS | -8.72, 34.10, 25.92 | 6.21, 31.04, 25.99 | -5.54, 25.16, 31.55 | 6.86, 26.19, 38.56 | -7.02, 36.94, -13.11 | 10.87, 35.06, -5.09 |
| P48CS | -3.34, 28.54, 24.20 | 5.87, 40.15, 17.96 | -7.58, 24.03, 37.93 | 11.91, 23.85, 40.98 | -8.06, 36.04, -6.05 | 13.66, 37.28, -17.09 |
| P47CS | -0.67, 25.84, 20.14 | 1.43, 25.49, 17.20 | -7.86, 12.20, 42.04 | 8.11, 11.45, 45.05 | -1.98, 40.24, -15.36 | 1.02,38.60,-13.11 |
| P43CS | -2.85, 23.88, 24.58 | 5.29, 27.07, 24.76 | -2.83, 9.40, 43.93 | 5.33, 8.85, 46.26 | -4.26, 38.19, -12.07 | 5.11,38.48,-16.47 |
| P41CS | 0.18, 26.56, 17.92 | 12.67, 27.14, 19.01 | -0.04, 12.88, 58.69 | 6.99, 15.53, 57.30 | -3.17, 37.28, -7.56 | 8.67, 38.04, -15.47 |

Table S3: MNI coordinates for microelectrode positioning in all patients in dACC, preSMA and vmPFC.

### Supplementary Figures

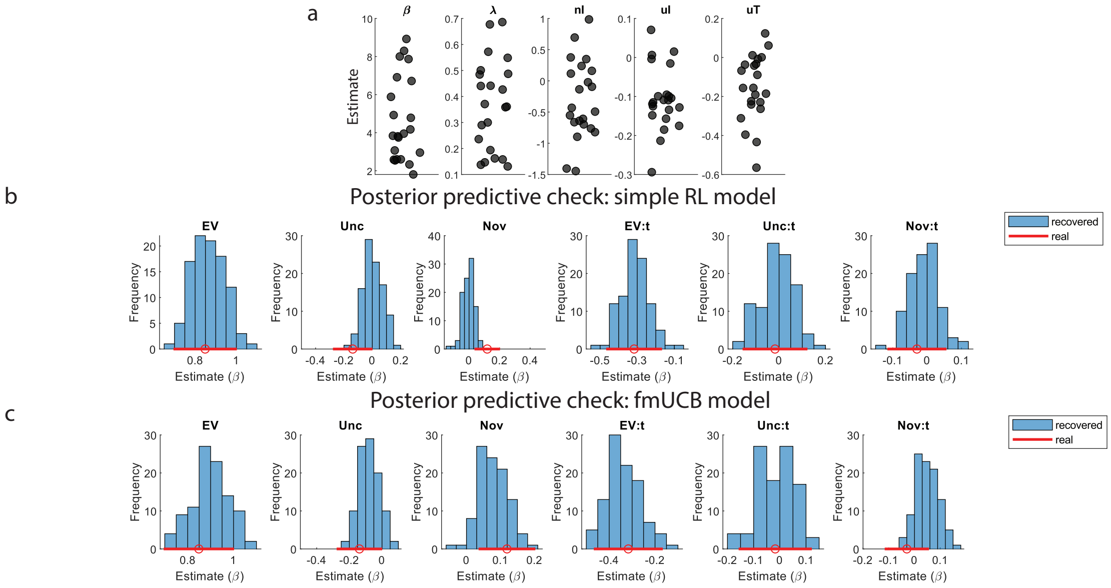


Figure S1: Model fits and posterior predictive check for selected exploration model with familiarity gating mechanism. (a) Individual fmUCB model parameter fits. Each dot represents a parameter fit for each patient (Left to right: softmax inverse temperature $\beta$; learning rate $\lambda$; novelty intercept; uncertainty intercept; uncertainty terminal). (b) Posterior predictive check for a simple reinforcement learning model which only included a softmax beta and a learning rate as free parameters. We fit this model to patient behavior and re-exposed an artificial agent with the obtained model parameters to the same set of trials which patients experienced 50 times, to generate decisions according to the estimated decision probabilities. We then fit a logistic regression for the effect of each variable (left to right: expected values, uncertainty, novelty, and their respective interactions with trial number) on decision in the artificial agents (blue histogram) and compared it to the actual estimate given true decisions concatenated across patients (red line; dot indicates mean and bars indicate 95% confidence interval). (c) Same, for the fmUCB model, which was selected for all subsequent analyses.

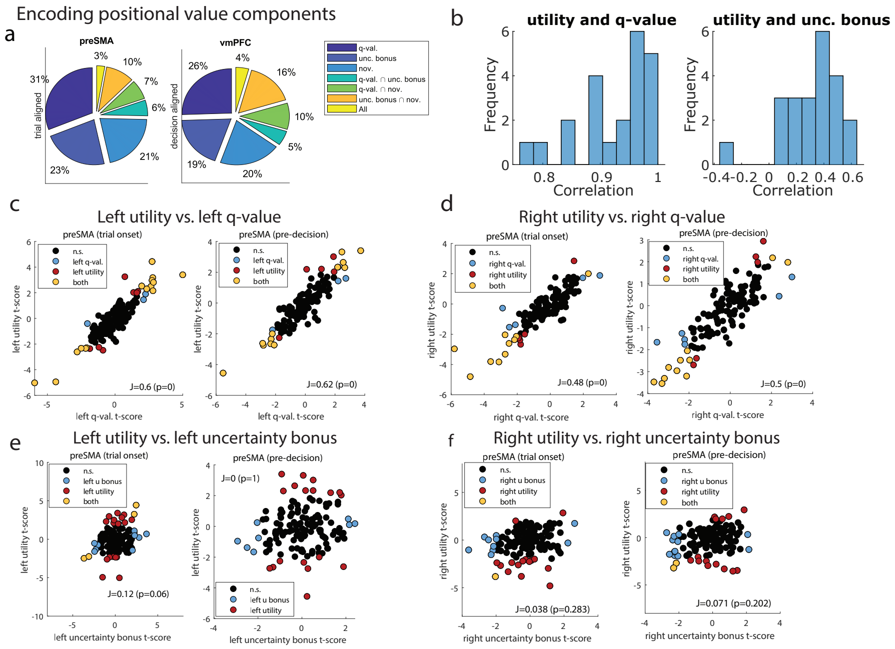


Figure S2: Summarizing positional encoding and comparing encoding of utility versus value components. (a) Pie chart including proportion of neurons sensitive to each positional component of value, and their respective overlaps, in preSMA (left) and vmPFC (right). (b) Histogram of correlation between utility and q-value (left), or utility and uncertainty bonus (right) trial vectors, across recording sessions. (c) Given the sizeable correlations between utility and its components, we mapped out the extent to which left utility preSMA neurons also correlated with left q-value preSMA neurons, in the trial onset period (left), and the pre-decision period (right). We plot q-value t-scores from the positional components GLM, as well as the utility t-scores from the utility and decision GLM (black: non-sensitive neurons; blue: left q-value neurons; red: left utility neurons; yellow: neurons concurrently sensitive for both). We tested whether this overlap is significant and report the Jaccard index (J), as well as p-values from the Jaccard test of overlap. (d) Same, for right utility vs. right q-value. (e) Same, for left utility vs left uncertainty bonus. (f) Same, for right utility vs. right uncertainty bonus.

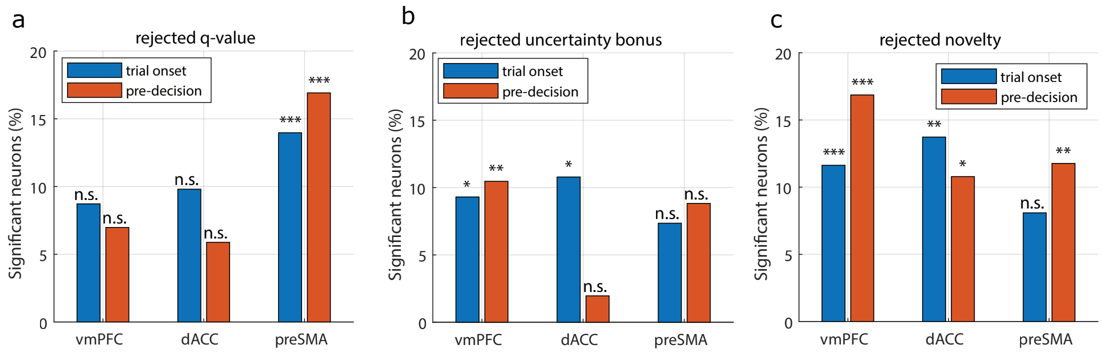


Figure S3: Single neuron encoding for the q-value, uncertainty bonus and novelty of the rejected stimulus in each trial. (a) Proportion of neurons sensitive to rejected q-value in vmPFC, dACC, and preSMA, in the trial onset (blue) and pre-decision (orange) periods. Stars indicate neuron count significance in a binomial test (* = p<0.05; ** = p<0.01; *** = p<0.001, Bonferroni corrected).

a

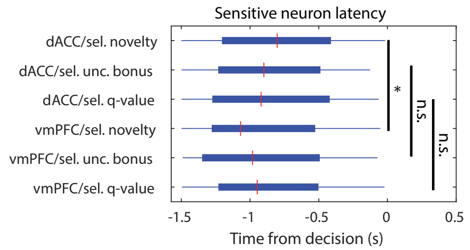


Figure S4: Timing for neurons which encode selected components of value. (a) Box plots of latency time across trials for all selected q-value, selected uncertainty bonus, or selected novelty neurons in vmPFC/dACC. The red mark indicates the median, and the box extends between the 25th and 75th percentiles of latency times. Bar whiskers extend to the most extreme data points not labeled as outliers, defined as values that are more than 1.5 times the interquartile length away from the edges of the box. Stars indicate significance in a two-sided rank-sum test between latencies for each regressor (* = p<0.05).

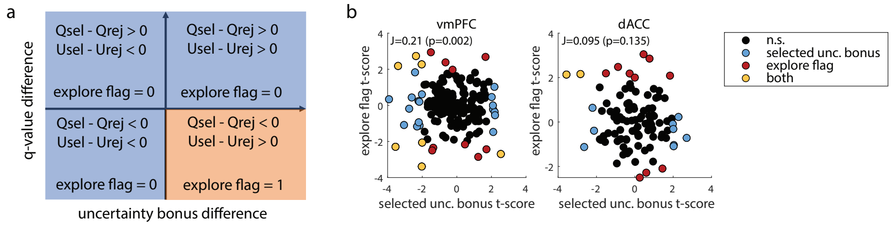


Figure S5: Comparing encoding of selected uncertainty bonus and exploration. (a) Chart indicating how trials were defined as explore or non-explore trials. Trials in which the selected option had lower q-value and higher uncertainty bonus were defined as explore trials (orange) and all other trials were defined as non-explore trials (blue). (b) Scatter plot of selected uncertainty t-scores versus explore flag t-scores from the Poisson GLM analysis. We display neurons sensitive to selected uncertainty bonus (blue), to the explore flag (red), and to both regressors (yellow). We also indicate the Jaccard overlap index and a p-value for the Jaccard test, indicating a significant overlap between uncertainty and exploration coding in individual vmPFC neurons. Left: vmPFC neurons; Right: dACC neurons.

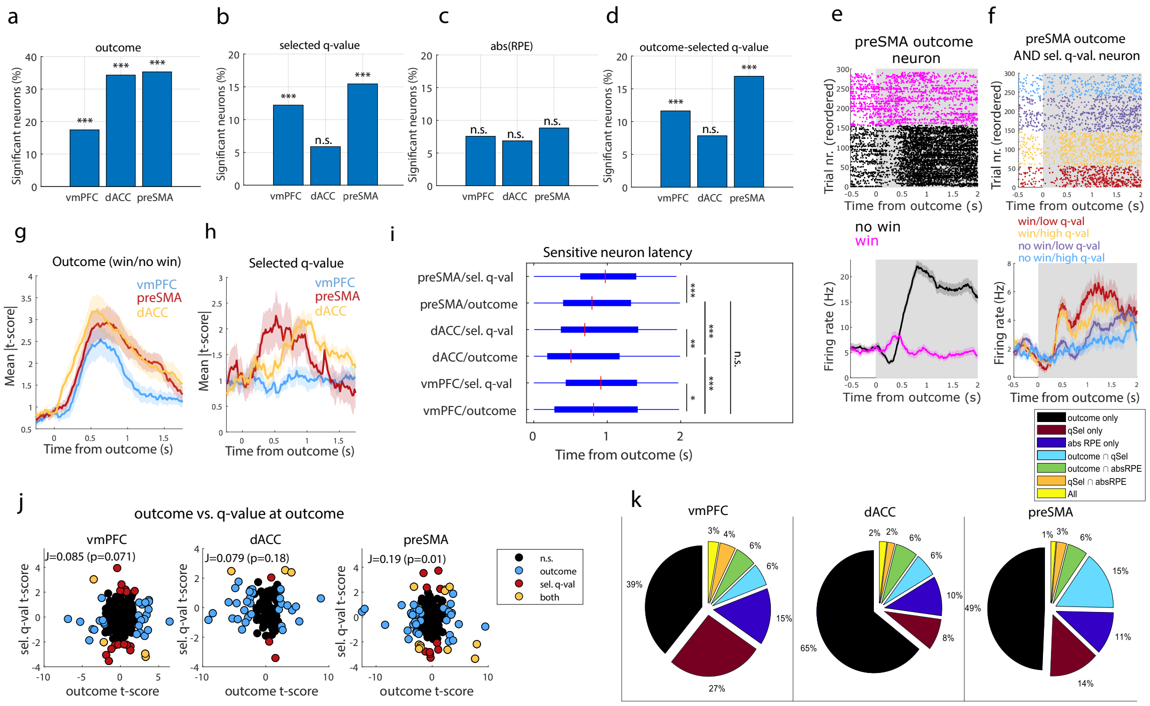


Figure S6: Post-feedback encoding. (a) Percentage of outcome neurons in vmPFC, dACC, and preSMA (*** = p<0.001, binomial test). (b) Same, for selected q-value. (c) Same, for abs(RPE). (d) Same, for the outcome minus selected q-value contrast. (e) Outcome neuron in preSMA. Top: Raster plots. For plotting, we sorted trials by outcome (magenta: win; black: no-win). Bottom: PSTH (bin size = 0.2 s, step size = 0.0625 s). Shaded areas indicate standard error. (f) Same, for an outcome and selected q-value preSMA neuron. Trials were split into outcome/q-value groups: win/low (red); win/high (yellow); no-win/low (purple); no-win/high (blue). (g) Mean absolute t-score in outcome neurons in vmPFC (blue), dACC (red), and preSMA (yellow). (h) Same, for selected q-value neurons. (i) Latency times box plot for outcome or selected q-value neurons in vmPFC, dACC, or preSMA. (* = p<0.05; ** = p<0.01; *** = p<0.001, two-sided rank-sum test). (j) Scatter plot of outcome versus selected q-value t-scores. We display neurons sensitive to outcome (blue), to selected q-value (red), or both (yellow). We indicate the Jaccard overlap index and a p-value for the Jaccard test. Left: vmPFC; Center: dACC; Right: preSMA. (k) Pie charts of neuron preference for outcomes, selected q-values, abs(RPE). Left: vmPFC; Center: dACC; Right: preSMA.
